# Act Natural: Functional Connectivity from Naturalistic Stimuli fMRI Outperforms Resting-State in Predicting Brain Activity

**DOI:** 10.1101/2021.11.01.466749

**Authors:** Shachar Gal, Yael Coldham, Niv Tik, Michal Bernstein-Eliav, Ido Tavor

## Abstract

The search for an ‘ideal’ approach to investigate the functional connections in the human brain is an ongoing challenge for the neuroscience community. While resting-state functional magnetic resonance imaging (fMRI) has been widely used to study individual functional connectivity patterns, recent work has highlighted the benefits of collecting functional connectivity data while participants are exposed to naturalistic stimuli, such as watching a movie or listening to a story. For example, functional connectivity data collected during movie-watching were shown to predict cognitive and emotional scores more accurately than resting-state-derived functional connectivity. We have previously reported a tight link between resting-state functional connectivity and task-derived neural activity, such that the former successfully predicts the latter. In the current work we use data from the Human Connectome Project to demonstrate that naturalistic-stimulus-derived functional connectivity predicts task-induced brain activation maps more accurately than resting-state-derived functional connectivity. We then show that activation maps predicted using naturalistic stimuli are better predictors of individual intelligence scores than activation maps predicted using resting-state. We additionally examine the influence of naturalistic-stimulus type on prediction accuracy. Our findings emphasize the potential of naturalistic stimuli as a promising alternative to resting-state fMRI for connectome-based predictive modelling of individual brain activity and cognitive traits.

## 1. Introduction

The field of cognitive neuroscience is ultimately aimed at understanding how our brains function in a real-life environment. How do we integrate the dynamic flow of multiple stimuli in a complex, multi-modal experience such as watching a play, taking a test, or meeting a friend? These real-life situations are quite challenging to simulate in laboratory settings. Naturalistic stimuli, such as listening to a story or watching a movie, are complex and dynamic stimulus which are used as ecologically valid conditions to measure neural activity (Eickhoff, Milham, & Vanderwal, 2020). While previous work has demonstrated a synchronized neural response across participants during movie-watching (Hasson, Malach, & Heeger, 2010; Hasson, Nir, Levy, Fuhrmann, & Malach, 2004), recent studies claim that the remaining individual variance in brain activity is more reflective of inherent, stable, connectivity features (Finn, 2021; Finn & Bandettini, 2021; Finn et al., 2017).

Still, functional connectivity data are most commonly collected using resting-state functional magnetic resonance imaging (rs-fMRI), by measuring the temporal correlations between spontaneous neural activation of different brain regions while participants are not performing any explicit task (van den Heuvel & Hulshoff Pol, 2010). Resting-state functional connectivity patterns are considered to represent an intrinsic functional organization of the human brain (Biswal, Zerrin Yetkin, Haughton, & Hyde, 1995; Greicius, Krasnow, Reiss, & Menon, 2003; van den Heuvel & Hulshoff Pol, 2010), and were shown to be closely associated with task-induced brain activity (Cole, Ito, Bassett, & Schultz, 2016; Parker Jones, Voets, Adcock, Stacey, & Jbabdi, 2017; Tavor et al., 2016; Tik et al., 2021). These studies have demonstrated that distinctive functional connectivity patterns are extremely successful predictors of task-evoked brain activity, in both healthy participants (Tavor et al., 2016) as well as psychiatric (Tik et al., 2021) and neurological patients (Parker Jones et al., 2017). Functional connectivity has also been reported to successfully predict individual scores of intelligence and cognition (Dhamala, Jamison, Jaywant, Dennis, & Kuceyeski, 2021; Dubois, Galdi, Paul, & Adolphs, 2018; Finn et al., 2015; Rosenberg et al., 2016; Song et al., 2008) as well as personality traits (Hsu, Rosenberg, Scheinost, Constable, & Chun, 2018; Nostro et al., 2018).

In the last few years, naturalistic stimuli derived data are increasingly being used to extract functional connectivity features (Finn et al., 2020; Vanderwal, Eilbott, & Castellanos, 2019) and seem to outperform rs-fMRI derived data in studying various neural and behavioral traits. For example, the use of naturalistic stimuli has been shown to enhance the detection of unique individual functional connectivity patterns compared to rs-fMRI, possibly due to increased arousal levels during movie-watching (Vanderwal et al., 2017). Functional connectivity data derived from movie-watching has also outperformed data collected at rest in predicting cognitive and emotional scores (Finn & Bandettini, 2021). Notably, the appropriate content to be used in naturalistic stimuli is debatable (Grall & Finn, 2021), with different studies employing different types of audiovisual content such as abstract videoclips as opposed to scenes from familiar films (Vanderwal, Kelly, Eilbott, Mayes, & Castellanos, 2015).

In light of previous evidence tightly linking functional connectivity patterns to task-induced brain activity, as well as evidence demonstrating the advantages of naturalistic stimuli compared to rs-fMRI, in the current work we examine the predictability of task-induced brain activity from patterns of brain connectivity while participants watch a naturalistic stimulus. We hypothesize that brain-connectivity data derived from movie-watching fMRI will be successful at predicting individual task-evoked neural activity across a range of cognitive paradigms. Moreover, we expect prediction of activation maps from naturalistic-stimulus data to outperform prediction from rs-fMRI data, and to successfully detect individually unique activation patterns. We additionally compare the accuracy of task-activation maps predicted from different types of naturalistic stimuli (Hollywood scenes (HO) vs. independent films made available under creative commons license (CC)). Finally, we demonstrate the predictability of cognitive scores from the predicted task-induced activation maps derived from functional connectivity data collected using a naturalistic stimulus, compared to functional connectivity data collected using rs-fMRI.

## 2. Methods

### 2.1. Participants

The data used in this study were provided by the Human Connectome Project (HCP; Van Essen et al., 2013). While the HCP dataset includes over 1000 participants, only a subset of 184 participants were scanned during naturalistic stimuli viewing, from which 175 participants were reported by the HCP to have good quality data. Naturalistic stimuli fMRI scans were performed on a 7 Tesla Siemens Magnetom. Since this work focuses on naturalistic stimuli, we used this subset of participants for whom naturalistic viewing data were available. Besides movie-watching scans, these participants were also scanned at rest on both 7T and 3T scanners, and during task performance on a 3T scanner. Participants that did not complete all movie-watching, resting-state and task fMRI scans were excluded from this study, resulting in a final dataset of n = 153 (mean age = 29.5, standard deviation = 3.24, 94 females). The HCP data includes multiple sibships of monozygotic and dizygotic twins as well as non-twin siblings. These family relations were accounted for in our analyses (see below).

### 2.2. Functional MRI data

#### 2.2.1. 7T data

The 7T dataset includes four resting-state scans and four movie-watching scans for each participant, acquired over four scanning sessions: Session #1 included two movie-watching scans and a resting-state scan; Sessions #2 and #3 included a resting-state scan only; Session #4 included two movie-watching scans and a resting-state scan. Therefore, the first two movie-watching scans were acquired in the same session as the first resting-state scan, while the third and fourth movie-watching scans were acquired in the same session as the fourth resting-state scan (see Table 1).

**Table 1.** Description of the movie-watching scans included in the 7T dataset.

| Movie run | Scanning session | Place in session | Stimuli type | Phase encoding |
| --- | --- | --- | --- | --- |
| 1 | 1 | 1 | Independent (CC) | AP |
| 2 | 1 | 2 | Hollywood (HO) | PA |
| 3 | 4 | 1 | Independent (CC) | PA |
| 4 | 4 | 2 | Hollywood (HO) | AP |

7T data were acquired using a gradient-echo-planar imaging (EPI) sequence with a TR of 1 second and a spatial resolution of 1.6 mm^3^. The phase encoding direction alternated between posterior-to-anterior (PA; Rest1, Movie2, Movie3) and anterior-to-posterior (AP; Rest4, Movie1, Movie4).

Each resting-state scan included 900 timepoints (15min long). Movie-watching scans varied slightly in duration between 901-921 timepoints, according to the clips being viewed during each scan. During movie-watching scans, participants were presented with a series of 4 or 5 audiovisual clips, separated by 20 seconds of rest. The first and third scans included clips from fiction and documentary independent films made available under creative commons license (CC), while the second and fourth scans included clips from Hollywood films (HO). Importantly, there was a dissociation between the phase-encoding direction of the scan and the type of movie presented in the scan (Table 1).

In order to retain the subsets of the data in which brain activity was affected by movie-watching, we excluded the last 17 seconds of each rest interval, keeping the first three seconds to account for the delay in the hemodynamic response. Following this exclusion procedure, the shortest scan consisted of 808 timepoints (Movie3). We then excluded timepoints from the end of each scan, both movie and rest, to keep a fixed number of 808 timepoints in all scans. Notably, while timepoints were excluded from movie-watching scans in an interleaved manner (i.e., from each no-movie interval), in the resting-state scans all timepoints were removed from the end of the scan. While this difference in timepoint-removal may have introduced a bias in favor of the resting-state data, as it made these data more continuous than the movie-watching data, it is noteworthy that such potential bias is in the opposite direction of our hypothesis.

In Addition, the HCP 7T dataset includes six retinotopy mapping scans for each participant (Benson et al., 2018) of which we used the four that have continuous visual stimulation (RETCCW, RETCW, RETCON, RETEXP) to examine the role of visual information in the prediction of task-activity maps. Each retinotopy scan included 300 timepoints. Similarly to the process described above, we removed the blank blocks (22 timepoints) from the beginning and end of each scan resulting in 256 timepoints per retinotopy scan. In order to compare prediction success across resting-state, movie-watching and retinotopy data, we then trimmed each movie-watching and resting-state scan to 256 timepoints (these shortened scans were used for the analysis presented in Figure S3 only, while for the main analysis we used the full-length resting-state and movie-watching data described above).

#### 2.2.2. 3T data

All functional 3T data were acquired with a TR of 0.72 seconds and a spatial resolution of 2 mm^3^. For the task-fMRI data, we used the z-score statistical maps provided by the HCP (see Barch et al., 2013) for a representative contrast from each task (see Table 2). While task-fMRI beta maps may be considered more “biological”, the z-score maps, computed as the beta divided by the noise variance, better address possible biases that may exist in the beta maps and were therefore chosen for our analysis. The Motor task was not included in the analysis due to the low inter-subject variability in its activation maps, resulting in low specificity of the predicted maps. For the resting-state fMRI, we used all four scans available and trimmed them similarly to the 7T resting-state scans, to include a total of 808 timepoints per scan.

**Table 2.** A list of the six fMRI-tasks included in this study. An exemplary contrast from each task in the HCP dataset, except the motor task, was used. See Barch et al., 2013 for more details regarding the tasks and their statistical analysis.

| Task | Contrast number | Contrast name |
| --- | --- | --- |
| Emotion | 01 | Faces-Baseline |
| Gambling | 02 | Reward-Baseline |
| Language | 02 | Story-Baseline |
| Relational | 02 | Rel-Baseline |
| Social | 02 | TOM-Baseline |
| Working Memory | 09 | 2BK-Baseline |

### 2.3. Preprocessing

We used the minimally preprocessed fMRI data provided by the HCP (Glasser et al., 2013). These data underwent motion and distortion corrections and nonlinear alignment to the MNI template space. Furthermore, the data were denoised using FMRIB’s ICA-based Xnoiseifier (FIX) (Griffanti et al., 2014; Salimi-Khorshidi et al., 2014), and then resampled and “projected” onto a surface representation of 91,282 vertices (“grayordinates”) in standard space. Data were aligned and registered using Multimodal Surface Matching (MSMAll) (Glasser et al., 2016; Robinson et al., 2014).

### 2.4. Predicting task-activation maps from task-free fMRI data

This work employs a previously developed method to predict task-induced brain activation maps from scans acquired while no explicit task is performed (Gal et al., 2021; Tavor et al., 2016; Tik et al., 2021). Here, we compared activation maps predicted from movie-watching scans, 7T resting-state scans and 3T resting-state scans, as well as the retinotopy scans. To create these maps (hereafter referred to as *connTask* maps), a set of 45 independent features was first extracted from each input type (i.e., movie-watching, 7T resting-state and 3T resting-state) separately. This was done by first defining a set of brain maps that can be used as seeds for creating connectivity maps. To do that on a group level, we first used an iterative principal component analysis (PCA; Smith et al., 2014) routine. We perform incremental PCA to an individual’s time series to reduce dimensionality, then iteratively concatenated subjects and run PCA again (keeping 1000 dimensions at each run). This low-memory-requirement procedure to combine large fMRI datasets was found to achieve similar accuracies as full temporal concatenations of all participants (Smith et al., 2014). We then applied group independent component analysis (ICA; Beckmann, DeLuca, Devlin, & Smith, 2005) on the concatenated data to yield a group-wise set of 45 components. These steps were performed on an independent set of 150 participants who were not included (and had no relatives) in further analyses, to prevent data leakage between the training and test sets. We then performed a dual regression (Beckmann, Mackay, Filippini, & Smith, 2009), resulting in 45 spatially independent cortical individual maps for each of our 153 participants, each map representing a brain network. These maps were then used as seeds in a weighted seed-to-vertex connectivity analysis. This process included a regression of these individual maps against individual time series to obtain an individual time course for each component, which were then correlated against the individual data to produce an individual spatial connectivity map per subject for each of the 45 components. The resulting connectivity maps were used as the final features for model fitting and prediction. Since our main interest in this work was to compare between different input types (i.e., naturalistic stimuli vs. rest) we deliberately did not perform modifications to the feature extraction steps but instead employed the exact pipeline as reported previously (Tavor et al., 2016). Figure 1 present a schematic illustration of the prediction procedure.

**Figure 1.**
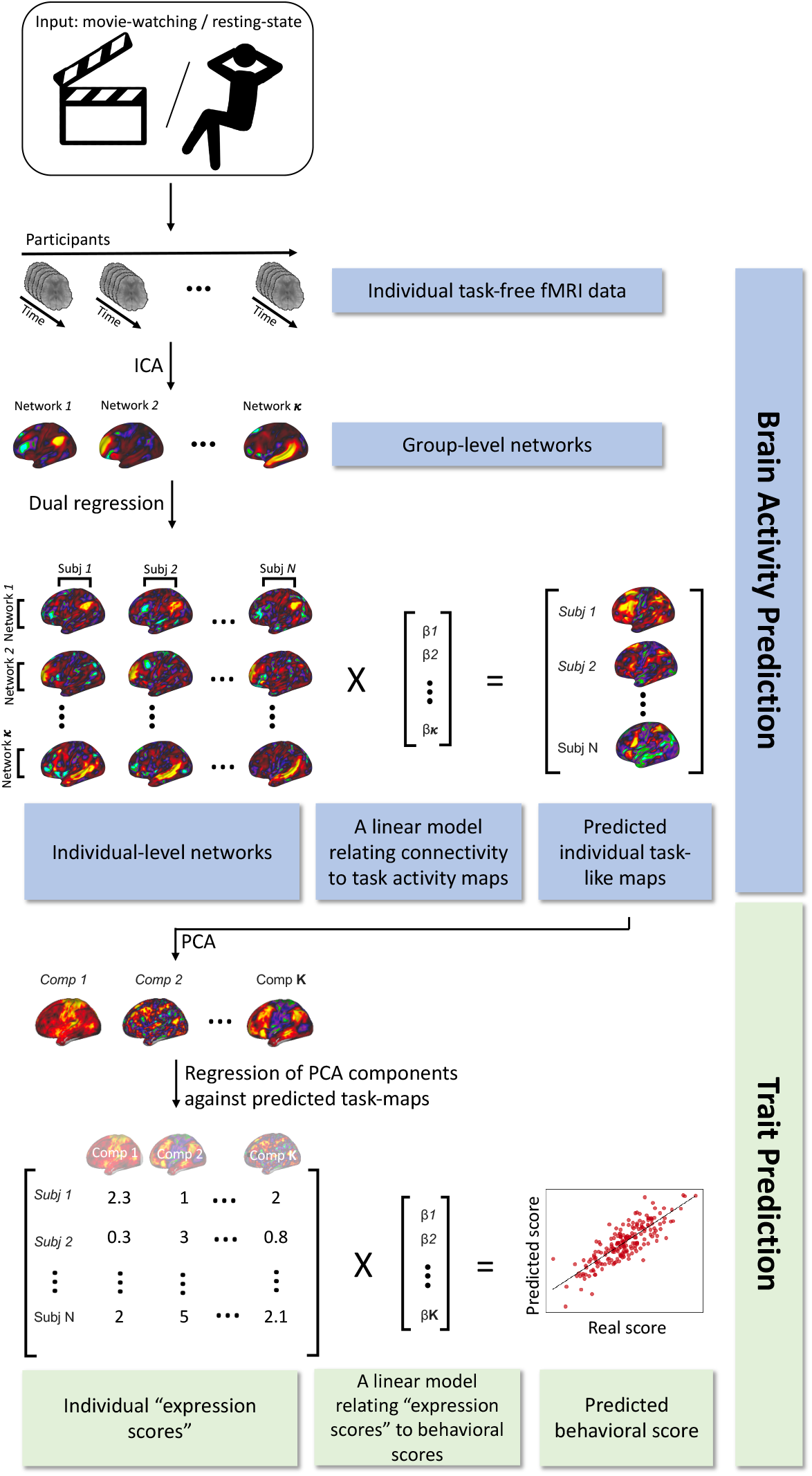
A schematic illustration of the prediction pipeline. Data from resting-state or movie-watching fMRI scans were first used to predict individual task-evoked brain activation maps (Blue). These predicted maps were then used to further predict individual cognitive abilities (Green).

These feature maps were then fed into a general linear model to predict task activation maps of a variety of tasks (Table 2). For each task, a 10-fold cross validation routine was employed to create a predicted *connTask* map, such that in each of the 10 iterations, 9/10 of the data were used as the training-set and 1/10 as the test-set. Taking into account the family structure in the HCP data, we made sure that in each iteration, members of the same family were kept in one group (either training or test) to prevent data leakage resulting from the heritability of connectivity patterns (Colclough et al., 2017).

Task-activity prediction was considered successful if the predicted map was more similar to the true activation map of the same participant than to all other participants’ maps (Gal et al., 2022; Tavor et al., 2016; Tik et al., 2021). In these cases, the correlation matrix between true and predicted maps is diagonal-dominant. To test for significance of this diagonality, we randomly shuffled this correlation matrix and compared the true accuracy of prediction (as measured by the diagonal mean) to 1000 permutated null diagonals.

### 2.5. Comparing the accuracy of task-activation maps predicted from different inputs

We tested whether *connTask* maps accuracy was higher when predicted from movie-watching data than from resting-state data. The accuracy of a *connTask* map was defined as the Pearson’s correlation coefficient between the predicted and actual activation maps. For each task-contrast, we performed a permutation test to determine whether the difference in accuracy between input types is statistically significant. To create a null distribution of the differences in prediction accuracy between movie-watching and resting-state, we randomly shuffled the prediction accuracy scores (i.e., the Pearson’s correlations between actual and predicted maps) between movie- and rest-derived maps and computed the difference between these scores. The p-value for significance was determined as the number of null differences that were equal or higher than the actual difference, divided by the number of permutations (5000). The resulting p values were corrected for 54 (18 contrasts X 3 input types) comparisons using Bonferroni correction for multiple comparisons.

Improved predictions of task-activity from movie-watching than resting-state may result from the visual information in the movies, considering that all tasks aside from the language task were visually presented. To examine the role of mere visual stimulation in task-activity prediction, we compared the accuracy of *connTask* maps derived from movie-watching, 7T resting-state and retinotopy data. This analysis was identical to the above analysis (comparing movie-watching, 7T resting-state and 3T resting-state) but employed the shortened movie-watching and resting-state scans, to match the length of the retinotopy scans.

Next, we explored the effect of audiovisual content type in the movie-watching scans on the accuracy of the *connTask* maps predicted from these scans. For this purpose, we generated *connTask* maps using data from each scanning session separately. For each participant and each scanning session, we defined a measurement of prediction improvement as the difference between the accuracies of the movie-watching-derived and the resting-state-derived *connTask* maps, using scans from the same session (i.e., movie scans 1&2 were compared with resting-state scan 1; movie scans 3&4 were compared with resting-state scan 4). We then grouped these improvement measures according to the movie-watching audiovisual content type (CC [1,3] or HO [2,4]), and performed a permutation test to determine whether the difference in accuracy between input types is statistically significant.. Resulting p values were corrected for six comparisons using Bonferroni correction for multiple comparisons.

Finally, we fitted a mixed linear model using ‘lme4’ package in R (Bates, Mächler, Bolker, & Walker, 2015) that explains the improvement in *connTask* maps accuracy by the audiovisual content (*stimuli_type*), with a random intercept across task contrasts, as well as a random intercept and slope across participants:

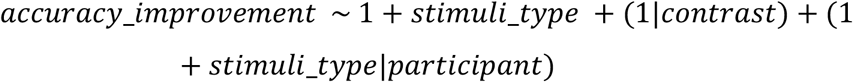

### 2.6. Relating the extent of improvement in accuracy to the level of brain activity

Improvement in prediction accuracy could arise from reduced predicted activity in regions where original activity was low (i.e., decreasing the “false positives”), or from increased predicted activity in regions where original activity was high (i.e., increasing the “true positives”). To examine what drives the effect of improved accuracy of movie-watching-derived task-activation maps relative to those derived from resting-state, we created maps that depict the mean activity and maps that depict the mean improvement of prediction by the movie-watching data, for 100 different functionally defined parcels. For this purpose we used a 100-node parcellation based on the division of the human cortex to seven distinct connectivity networks (Schaefer et al., 2018; Yeo et al., 2011).

The mean activation was calculated as the average z-score across voxels within each parcel, for each task-contrast. Prediction improvement was calculated for each participant and each session as the difference between the accuracies of the movie-watching-derived and the resting-state-derived *connTask* maps, using scans from the same session. We then averaged prediction improvement values across participants and sessions to create the parcellated maps. We created such parcellated maps for each task-contrast and examined the correlations between prediction improvement and mean activity in each task separately.

To further examine the relationship between prediction improvement and task-activity beyond the level of a single task, we used a mixed linear model (Bates et al., 2015) that explains the improvement in *connTask* accuracy in each parcel by the mean activity in this parcel in the original task-activation map, with a random intercept and slope across task contrasts and across parcels:

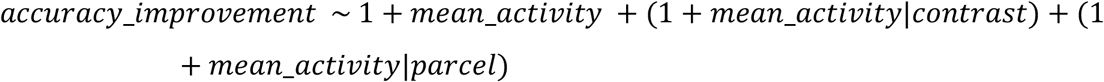

Notably, the degrees of freedom in this model are limited by the number of task-contrasts we used. In this case, only six task-contrasts do not allow sufficient data for fitting a reliable model. Therefore, we used twelve more task-contrasts from the HCP dataset, as in previous work on *connTask* maps (Gal et al., 2022).

### 2.7. Prediction of individual traits using *connTask* maps

While it has been previously shown that connectomes derived from movie-watching data outperformed those derived from resting-state data in the prediction of individual traits (Finn & Bandettini, 2021), here we tested whether a similar effect also exists when using *connTask* maps rather than the traditional connectome.

We used the resting-state and movie-watching-derived *connTask* maps to predict individual scores of General Cognitive Ability (GCA). GCA scores were constructed using an exploratory factor analysis, following the procedure suggested by Dubois et al. (2018) and the code made available by them. We used unadjusted scores from ten cognitive tasks, including seven tasks from the NIH Toolbox (Dimensional Change Cart Sort, Flanker Task, List Sort Test, Picture Sequence Test, Picture Vocabulary Test, Pattern Completion Test, Oral Reading Recognition Test), and three tasks from the Penn Neurocognitive Battery (Penn Progressive Matrices, Penn Word Memory Test, Variable Short Penn Line Orientation Test). The G-scores prediction was performed using the Basis Brain Set pipeline (BBS) (Sripada et al., 2019; Sripada, Angstadt, Rutherford, Taxali, & Shedden, 2020), which has been previously found effective for predicting individual traits from *connTask* maps (Gal et al., 2022) (Figure 1).

We performed a 5-fold cross validation routine while taking into account the family structure in the HCP data. In each iteration, a model was trained on 4/5 of the data, and predictions were yielded for 1/5 of the data, while genetically related participants kept in either the training or the test datasets. For each input type (i.e., resting-state or movie-watching) and each task, the prediction of GCA scores from *connTask* maps included the following steps: (1) dimensionality of the training set data was reduced using PCA to a predetermined number of components, k. We used k=20 to maintain a similar ratio between the number of observations (participants) and the number of predictors (components) as in previous work (Gal et al., 2022). (2) These components were regressed against the individual data of each participant, test and training set alike, to yield an “expression score” for each component. (3) The training sets’ expression scores were used to fit a linear model that predicts the GCA. (4) The model was applied on the test set. For significance testing, this procedure was repeated 1000 times with 2/3 of the data used for training and 1/3 for testing, accounting for family structure. Differences between the accuracy of predictions based on the two input types were tested using a non-parametric, related samples test (Wilcoxon signed ranks test) (Demšar, 2006).

## 3. Results

### 3.1. Movie-watching-derived *connTask* maps are more accurate than resting-state-derived maps

All predictions of task-activation maps, regardless of the input type, were statistically significant (*p* < 0.005), as evaluated by a permutation test (see Methods). Task-activation maps predicted from movie-watching data were significantly more accurate than those predicted from 7T resting-state data for all task-contrasts (Figures 2, 3 and Supplementary Figures S1, S2). Additionally, no significant differences were observed between the accuracy of *connTask* maps derived from 3T and 7T resting-state data, demonstrating that the strength of magnetic field does not affect prediction accuracy, while *connTask* maps derived from movie-watching data outperformed both. Task-activation maps predicted from retinotopy data were significantly less accurate than those predicted from movie-watching data across all task-contrasts (Supplementary Figure S3). For 9 out of 18 task-contrasts, retinotopy outperformed the resting-state data in predicting task-activation maps.

**Figure 2.**
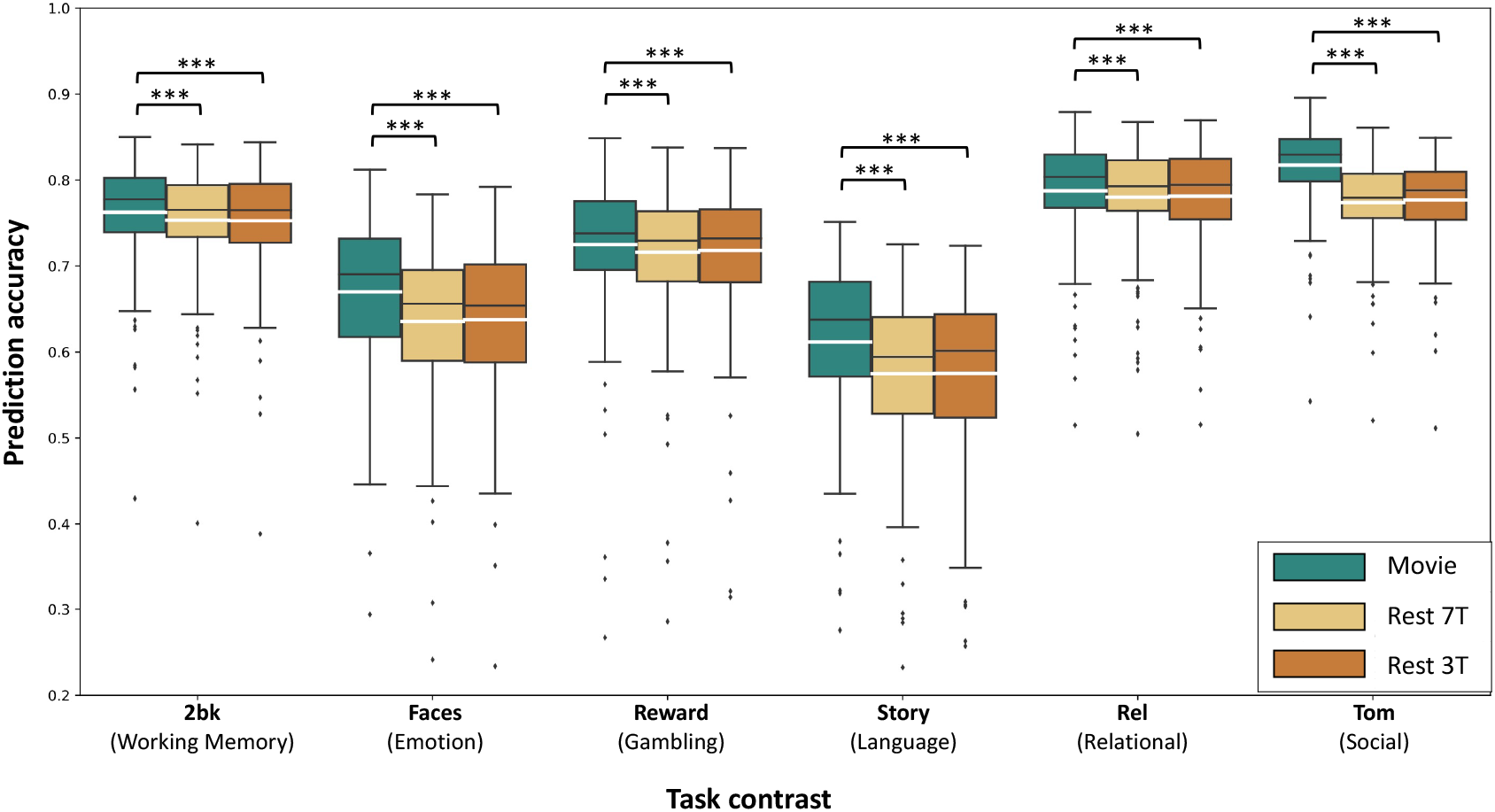
Movie-watching-derived *connTask* maps are more accurate than resting-state-derived maps. For each task-contrast and each input type (movie-watching, 3T resting-state and 7T resting-state), boxplots portray the distribution of prediction accuracies. Prediction accuracy for each participant was calculated as the Pearson correlation between *connTask* and actual task-induced activation maps. Black lines represent the median, white lines represent the mean. Differences in *connTask* accuracy between input types and within each task-contrast were tested using a permutation test and corrected using Bonferroni correction for multiple comparisons. \*\*\**p* < 0.001

**Figure 3.**
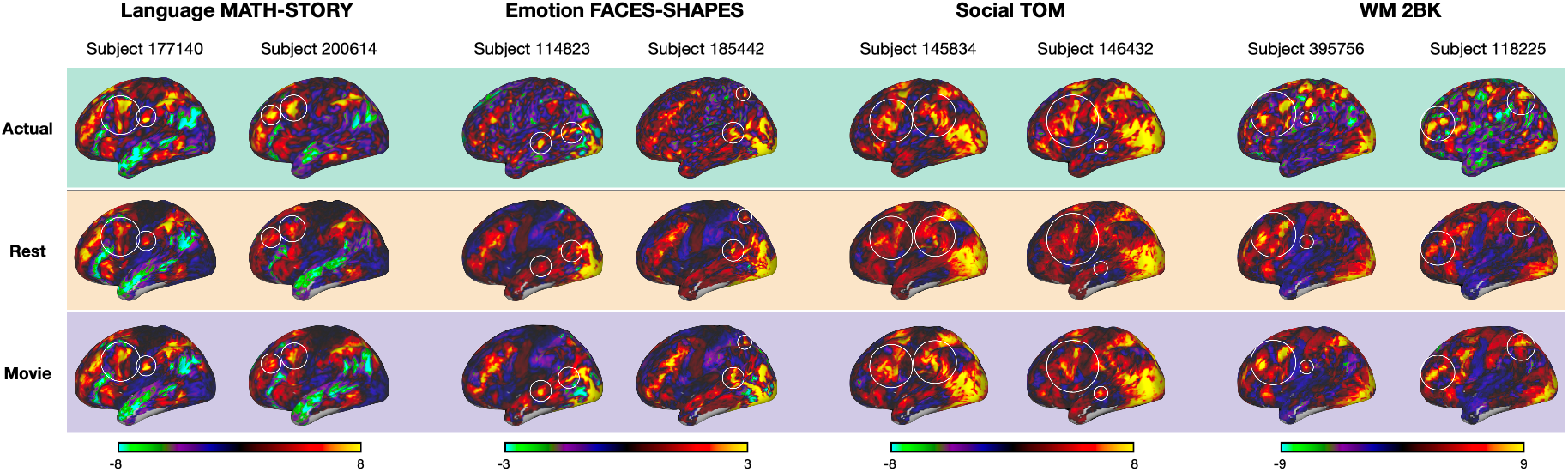
Examples of actual brain activation maps (top) and the corresponding maps predicted from either resting-state (middle) or movie-watching fMRI (bottom). Examples are presented for 4 different task-contrasts. Brain activation maps are depicted on the surface representation of the cortex, white circles highlight areas of accurate predictions of individual-specific activation patterns.

### 3.2. The type of naturalistic stimulus impacts the accuracy of the predicted task-activation maps

In four of the six task-contrasts, we found that *connTask* maps predicted from movie-watching data where participants viewed independent films (CC) compared to Hollywood films (HO) offered a larger improvement in the prediction of activation patterns, relative to the prediction from resting-state data (Figure 4 and Supplementary Figure S4). This effect was observed in 7 out of 12 additional task-contrasts, with moderate to large effect sizes (Supplementary Table S1).

**Figure 4.**
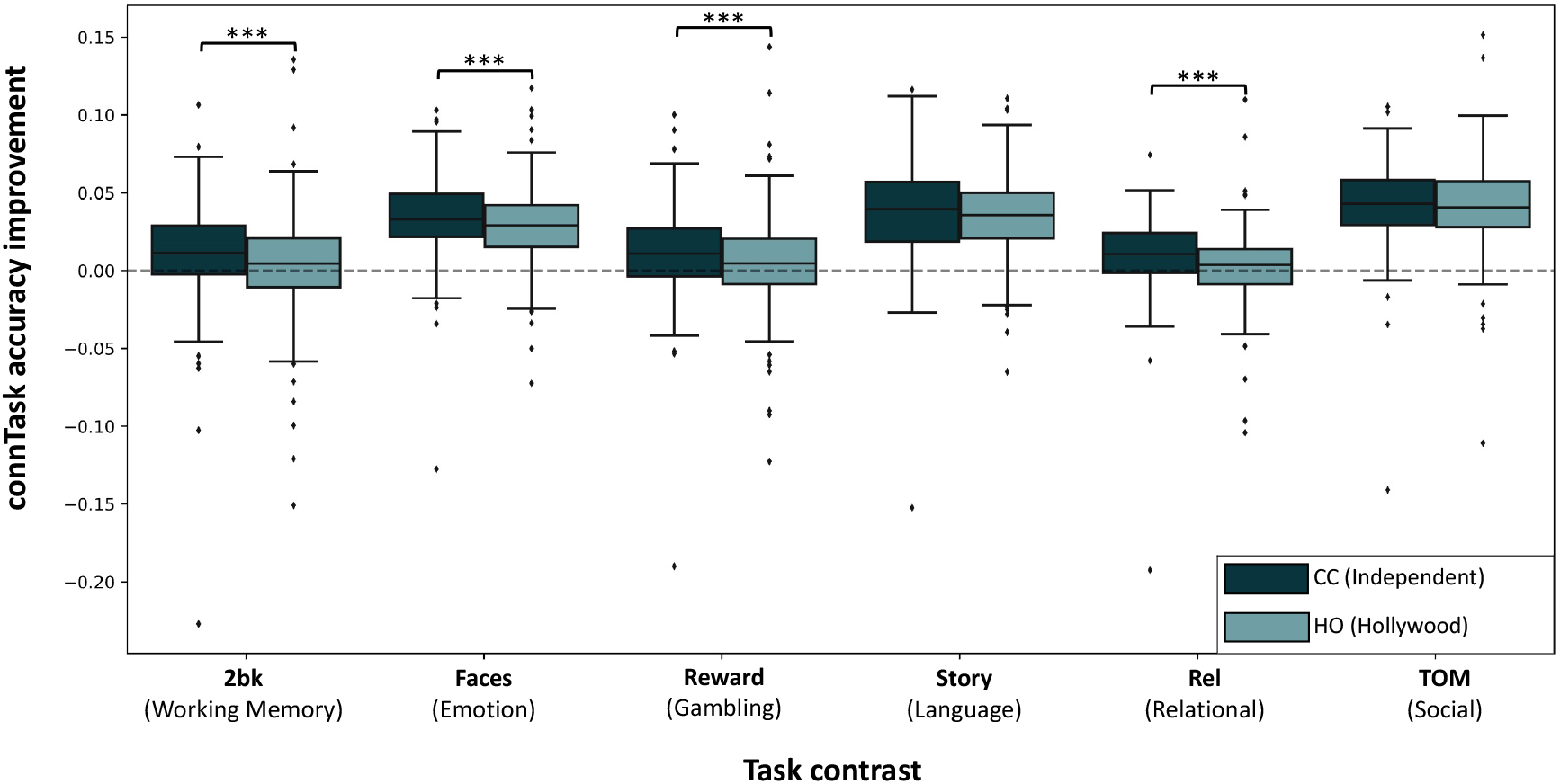
The type of naturalistic stimuli impacts the accuracy of subsequent *connTask* map. For each task-contrast and each type of audiovisual content, boxplots portray the distribution of prediction improvement. For each participant and each content type, we defined prediction improvement as the difference between the accuracies of the movie-watching-derived and the resting-state-derived *connTask* maps, using scans from the same session. Differences in accuracy improvement between input types and within each task-contrast were tested using a permutation test and corrected using Bonferroni correction for multiple comparisons. \*\*\**p* < 0.001

A mixed linear model that explains improvement in *connTask* accuracy by the input type while controlling for variance driven by the different tasks and participants further supported this effect (*p < 0.001, df=152)*, suggesting that task-activation maps predicted from data of CC naturalistic stimuli were more accurate than those predicted from HO naturalistic stimuli.

To examine whether differences in prediction accuracy between resting-state and movie-watching and between movie types may result from differences in head motion during these scans, we compared the mean framewise displacement between resting-state and movie-watching runs within each session. No consistent differences in head motion were observed (Supplementary Figure S5).

### 3.3. Improvement in task-activity prediction from movie-watching data is larger in highly active brain areas

In four of the six task-contrasts, prediction improvement was significantly correlated with the absolute values of mean activation across parcels (Figure 5). This effect was observed in 10 out of 12 additional task-contrasts and corroborated by the results of the mixed linear model analysis, showing a significant relationship between mean activity and prediction improvement across task-contrasts and brain areas (*p = 0.01, df=17)*.

**Figure 5.**
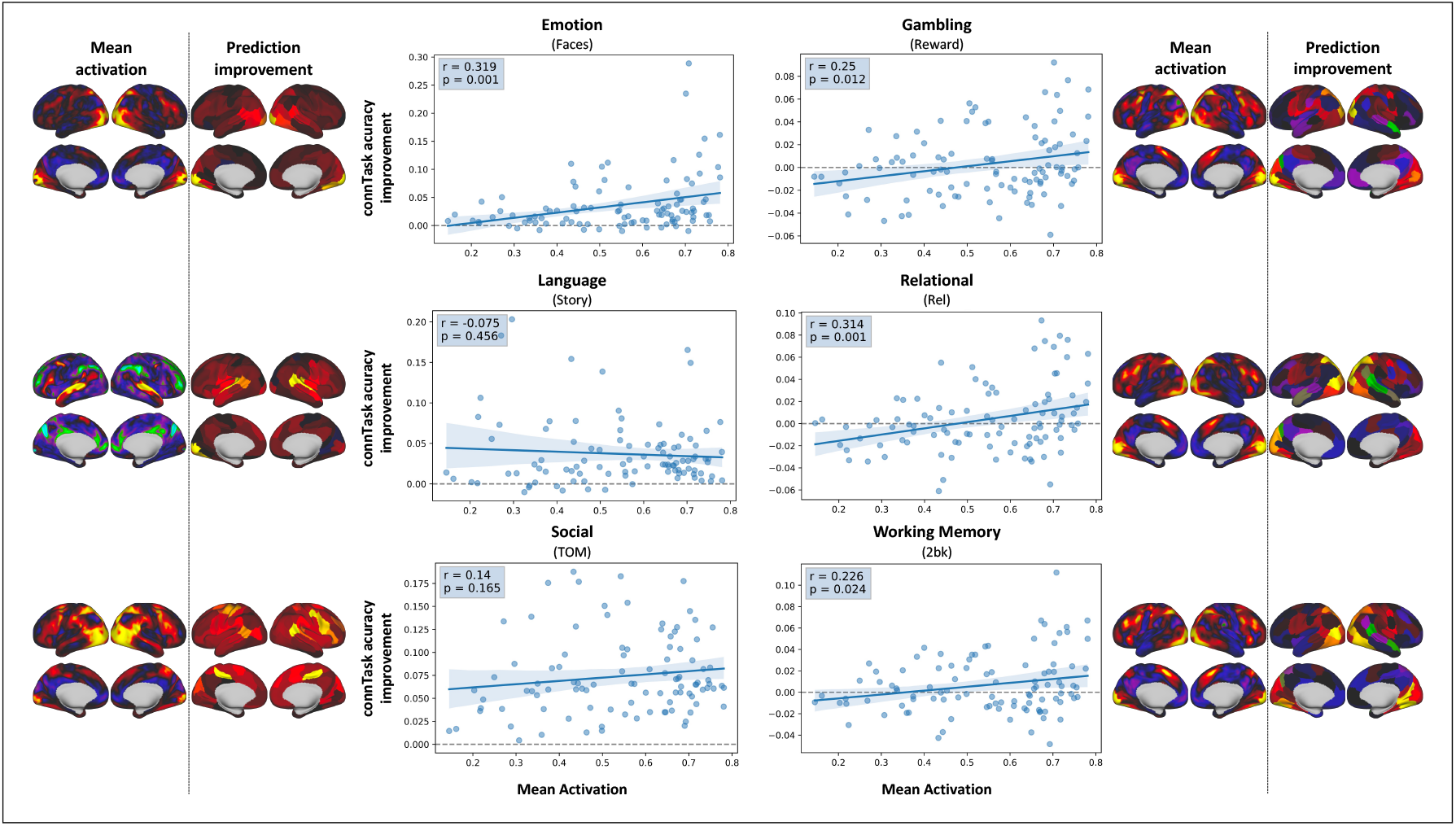
Improvement in task-activity prediction from movie-watching data is larger in highly active brain areas. Center: Scatter plots depicting the relationship between mean activation and prediction improvement in 100 distinct functionally defined parcels (Schaefer et al., 2018). Left and right: mean activation maps and parcellated prediction improvement maps for each contrast.

### 3.4. Prediction of individual traits from movie-watching-derived *connTask* maps is more accurate than from resting-state-derived maps

Individual scores of General Cognitive Ability (GCA) were successfully predicted from *connTask* maps derived from both movie-watching and resting-state data. All models predicted significantly better than chance, as confirmed by a permutation test with 5000 iterations (p<0.05, Bonferroni corrected for multiple comparisons). Models based on movie-watching-derived *connTask* maps were significantly more accurate than models based on resting-state-derived *connTask* maps for all 18 task-contrasts except the Social Random vs. TOM contrast (p<0.0001, Bonferroni corrected for multiple comparisons).

## 4. Discussion

Functional connectivity patterns derived from data collected while participants were exposed to naturalistic stimuli outperformed connectivity patterns derived from resting-state fMRI in predicting inherent individual qualities, including task-induced brain activity and cognitive scores. While it has been previously demonstrated that data collected at rest can be used to successfully predict brain activity during task performance (Cole et al., 2016; Parker Jones et al., 2017; Robinson et al., 2014; Tavor et al., 2016; Tik et al., 2021), here we demonstrate that data collected during movie-watching yield an even more accurate prediction of task-derived activation maps, across six different tasks: Emotion, Gambling, Language, Relational, Social and Working Memory (Table 2; Figures 2, 3). Moreover, we found that connectivity patterns derived from data acquired while watching independent films (CC) provided higher accuracy of the predicted task-activation maps than Hollywood scenes (HO) in most task-contrasts (Figure 4). Finally, task-activation maps derived from movie-watching outperformed those derived from rest in predicting individual cognitive scores (Figure 6).

**Figure 6.**
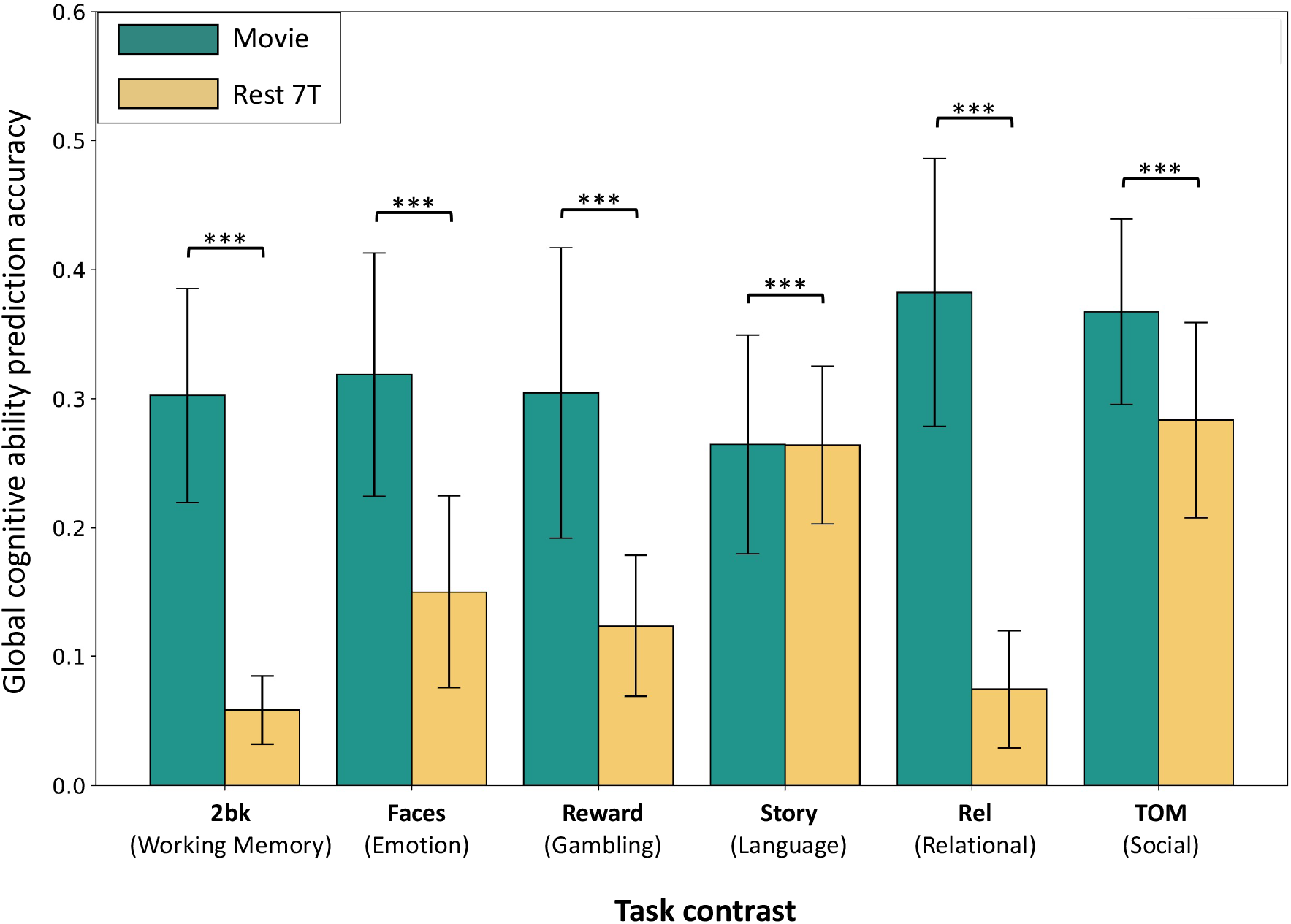
Prediction of individual traits is more accurate from movie-watching than resting-state-derived *connTask* maps. For each task-contrast and input type, bars height depicts the accuracy of GCA scores prediction, calculated as the Pearson correlation between real and predicted GCA scores. Error bars depict the standard error of prediction accuracy across the 5 folds of cross-validation. While all models predicted significantly more accurate than the null hypothesis (p<0.05, Bonferroni corrected for multiple comparisons), models based on movie-watching-derived *connTask* maps were significantly more accurate than models based on resting-state-derived *connTask* maps (p<0.0001, Bonferroni corrected for multiple comparisons).

The ability to predict task-evoked brain activation from task-free fMRI is of immense value. It provides a method for inferring multiple individualized functional localizers based on a single scan, without the need to acquire often time-consuming and cognitively demanding fMRI tasks. Thus, it could be used to investigate functional regions in patients who are unable to perform tasks, such as cognitively impaired, paralyzed, or pediatric patients. While previously demonstrated with resting-state fMRI data (Tavor et al., 2016), it remains an open question whether resting-state is indeed the optimal task-free paradigm for making such predictions. Recently, it has been suggested that the use of naturalistic stimuli may accelerate progress in areas that have been so far dominated by rest, including characterizing brain functional organization and brain–behavior relationships (Finn, 2021).Our results are in line with previous studies showing that movie-watching data outperform resting-state data in detecting unique functional connectivity patterns (Vanderwal et al., 2017) and that the prediction of cognitive and emotional scores is more accurate when based on movie-watching data than on data collected at rest (Finn & Bandettini, 2021). The superiority of movie-watching over rest may be attributed to the fact that movie-watching is a dynamic, ecologically valid stimulus, designed to resemble real-life experiences while measuring neural activity (Eickhoff et al., 2020). As such, movie-watching has been associated with increased participant compliance in the form of decreased head motion and sleepiness (Sonkusare, Breakspear, & Guo, 2019; Vanderwal et al., 2017, 2015) and with higher test-retest reliability (Wang et al., 2017) and less susceptibility to influences of volatile factors such as mood variations, level of concentration and tiredness (Christoff, Gordon, Smallwood, Smith, & Schooler, 2009; Duncan & Northoff, 2013; Finn et al., 2017; Harrison et al., 2008; Morcom & Fletcher, 2007; Shirer, Ryali, Rykhlevskaia, Menon, & Greicius, 2012).

We compared the mean framewise displacement between resting-state and movie-watching scans to examine whether difference in head motion may account for the superiority of movie-watching over rest in predicting brain activity. No consistent differences in head motion were observed (Supplementary Figure S5), suggesting that motion is unlikely to be the cause of differences in prediction accuracy between input types.

The improvement in task-activity predictions from movie-watching relative to resting-state data may also be explained by the visual information that is available in the movies, considering that all of the predicted tasks aside from the language task were based on visual presentation. To examine the role of visual information in task-activity prediction, we used the HCP’s 7T retinotopy scans, in which participants were presented with visual stimulation but with no naturalistic content. We found that predictions based on retinotopy data were significantly less accurate than those derived from movie-watching data (Supplementary Figure S3), suggesting that the mere visual stimulation is not enough, but rather the actual naturalistic content in the movies is what drives the improved prediction. Interestingly, for 9 out of 18 task-contrasts retinotopy data yielded more accurate predictions relative to resting-state data, suggesting that some visual stimulation may be better than none, but the effect of the exact type and content of stimulation should be further investigated. In efforts to improve the prediction of human behavior and brain activity from functional connectivity, most research to date focuses on model enhancement. This includes exploring different machine-learning algorithms and testing different model parameters (Ngo, Khosla, Jamison, Kuceyeski, & Sabuncu, 2021; Zheng et al., 2021). Here, we take an alternative data-centric approach and focus on enhancing the features rather than the model. As such, by feeding the model with connectivity features derived from movie-watching fMRI rather than resting-state we were able to significantly improve model performance.

Our findings provide important insights into the relationship between task-evoked activations and brain activity under naturalistic conditions. Movie-watching may elicit a cognitive state in which functional brain networks are organized similarly to their organization under task conditions, making it optimal for obtaining multiple task-activity maps from a single fMRI scan. The fact that task maps are better predicted from movie-watching than rest suggests that naturalistic viewing may be a better proxy for the cognitive mechanisms that operate during task performance, while still being short and effortless as opposed to actual tasks. Watching a few minutes of such rich, dynamic stimuli may utilize similar functional mechanisms to those that are revealed in traditional task-based fMRI.

The ‘ideal’ type of naturalistic stimuli to be used for predictive models is an open question (Grall & Finn, 2021). Here, we found that CC stimuli (independent films) offered a larger improvement in prediction of task-activity than HO stimuli (Hollywood films) relative to rest. This result was consistent across the Emotion, Gambling, Relational, Social and Working Memory tasks, but not the Language and Social tasks (Figure 4). Notably, these two task-contrasts demonstrated the highest improvement in prediction accuracy to begin with (Figure 2), alluding to a possible ceiling effect.

While CC videoclips outperformed HO videoclips in the prediction of some task-induced activation maps, previous studies have suggested that different types of naturalistic stimuli could be beneficial for different purposes. For example, in a recent work that used the HCP data in a connectome predictive modelling (CPM) paradigm, HO videoclips with high social content provided better predictions of cognitive scores than CC videoclips, with low or no social content (Finn & Bandettini, 2021). Notably, this result diverges from ours in the matter of which naturalistic stimulus yields superior predictions, emphasizing the variability offered by different stimuli in different prediction schemes. Moreover, different types of naturalistic stimuli were found to evoke different patterns of brain activation, with spatially limited and weaker synchronized activity across participants (Hasson et al., 2004) when watching videoclips of abstract shapes with no narrative compared to Hollywood movie clips (*Ocean’s Eleven*) (Vanderwal et al. 2015). Taken together, these findings highlight the crucial impact of the choice of stimulus on the obtained results. Hence, future studies based on data collected while participants are exposed to naturalistic stimuli should thoroughly consider the type of stimulus to be used.

The improvement in prediction accuracy of task-activation maps reported here for movie-watching relative to resting-state data could be explained by either reduced predicted activity in regions where original activity was low (i.e., decreasing the “false positives”), or by increased predicted activity in regions where original activity was high (i.e., increasing the “true positives”). We revealed a positive correlation between the original task-induced activation levels and the improvement in activation prediction from movie-watching compared to resting-state data, suggesting that the improvement in prediction was due to enhanced prediction of actual activation patterns, rather than reduced prediction in areas of low actual activity. This result was consistent across 14 out of 18 task-contrasts, excluding the language (story-baseline; math-baseline) and social (TOM-baseline; random-baseline) contrasts. However, the contrasts between two conditions in these two tasks (math-story and TOM-random) did show a significant effect. It is possible that the shapes stimuli presented in the social task, when contrasted against baseline, evoke relatively low-level visual activations that do not correspond with the improvement in prediction, but when contrasted with each other they evoke higher-level socially related brain activity. As for the language task, the auditory nature of the stimuli may account for this effect.

Previous research has shown that activity maps derived from resting-state data better predict cognitive scores than traditional functional connectivity properties (i.e., the resting-state connectome) (Gal et al., 2022). Here, we show that activity maps derived from data collected while viewing naturalistic stimuli outperform those that were derived from data collected at rest in the prediction of general intelligence scores (Figure 6). This finding may result from the fact that movie-watching data outperform resting-state data in the prediction of task-derived activation, however it may also arise from the former being a better predictor of intelligence than the latter, independently from task-activity prediction. The question of whether the increased accuracy is due to the better activity prediction or due to the suitability of movie-watching for predicting intelligence should be addressed in future studies exploring the uses and benefits of naturalistic stimuli. Regardless, this finding suggests that naturalistic stimuli designed to resemble complex, multi-modal real-life situations, may be more appropriate for the prediction of complex cognitive constructs.

Several limitations of the current work should be noted. First, the HCP dataset includes movie-watching scans acquired in 7T, resting-state scans acquired in 3T as well as 7T, and task-fMRI scans acquired in 3T. As such, the magnetic field strength differed between the data we used as the predictor (movie-watching) and the data we aimed to predict (task-fMRI). To examine whether the strength of the magnetic field may have introduced a bias to our analysis, we compared the accuracy of activation maps predicted from 7T rest data and 3T rest data. As exact T2* effects of individual participants’ anatomy depend on the field strength, one may expect that predictions of task-activity maps (that were acquired in 3T) would be more accurate from 3T than 7T resting-state data. Nevertheless, no significant differences were found in the accuracy of prediction based on 7T rest data and 3T rest data. We therefore conclude that the use of data acquired at 7T to predict activation maps could not have biased our results in favor of movie-watching over resting-state data.

Second, this study was performed on 153 HCP participants who have completed all movie-watching, resting-state, and task fMRI scans. While it has been suggested that associations between neural and behavioral measures only stabilize and become reproducible with sample sizes of n ⪆ 2,000 (Marek et al., 2020), a recent work that examined the required sample size for the prediction of task-induced activity from functional connectivity has shown that a training set of 150-200 participants should be sufficient for accurate predictions (Cohen, Chen, Jones, Niu, & Wang, 2020).

## 5. Conclusions

The findings reported in this work highlight the benefits of using functional connectivity derived from naturalistic-stimuli rather than resting-state fMRI for the prediction of individual task-induced brain activity and cognitive traits. Our results also emphasize the influence of the type of naturalistic stimulus on prediction accuracy, suggesting that further research is required in order to standardize and determine the ‘optimal’ naturalistic stimulus, if exists. We conclude that naturalistic stimuli such as movie-watching, that are designed to resemble complex, real-life experience, are a promising approach for unravelling the intrinsic functional organization of the human brain.

## CRediT authorship contribution statement

**Shachar Gal:** Conceptualization, Software, Formal analysis, Visualization, Writing - Original Draft. **Yael Coldham**: Writing - Original Draft. **Niv Tik:** Writing – Review & Editing, Formal analysis. **Michal Bernstein-Eliav**: Writing - Review & Editing. **Ido Tavor:** Conceptualization, Software, Supervision, Writing - Review & Editing

## Acknowledgments

Data were provided by the Human Connectome Project, WU-Minn Consortium (Principal Investigators: David Van Essen and Kamil Ugurbil; 1U54MH091657) funded by the 16 NIH Institutes and Centers that support the NIH Blueprint for Neuroscience Research; and by the McDonnell Center for Systems Neuroscience at Washington University.

The authors acknowledge with thanks the support of the Israel Science Foundation (ISF grant no. 1603/18) and the National Institute of Psychobiology in Israel (grant 232-19-20.).

## Data and materials availability

All the data included in this study are publicly available in the Human Connectome Project database at https://db.humanconnectome.org

