## Supplementary for "Act Natural: Functional Connectivity from Naturalistic Stimuli fMRI Outperforms Resting-State in Predicting Brain Activity"

### Supplementary Information

Shachar Gal[1,2], Yael Coldham[1,2], Niv Tik[1,2], Michal Bernstein-Eliav[1], Ido Tavor[1,2,3]

[1]Sackler Faculty of Medicine, Tel Aviv University, Tel Aviv, Israel

[2]Sagol School of Neuroscience, Tel Aviv University, Tel Aviv, Israel

[3]Strauss Center for Computational Neuroimaging, Tel Aviv University, Tel Aviv, Israel

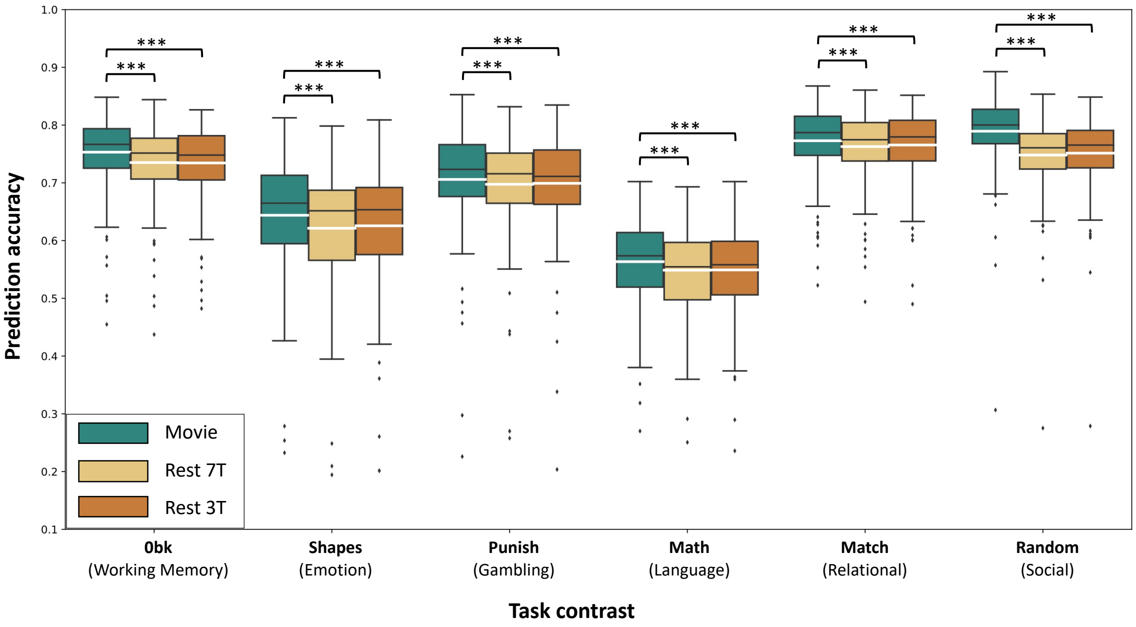


**Supplementary Figure S1. Movie-watching-derived *connTask* maps are more accurate than resting-state-derived maps.** This figure is similar to Figure 2 in the main test but shows additional task-contrasts of a single condition vs. baseline. For each task-contrast and each input type (movie-watching, 3T resting-state and 7T resting-state), boxplots portray the distribution of prediction accuracies. Prediction accuracy for each participant was calculated as the Pearson correlation between *connTask* and actual task-induced activation maps. Black lines represent the median, white lines represent the mean. Differences in *connTask* accuracy between input types and within each task-contrast were tested using a permutation test and corrected using Bonferroni correction for multiple comparisons. $^{***}p < 0.001$

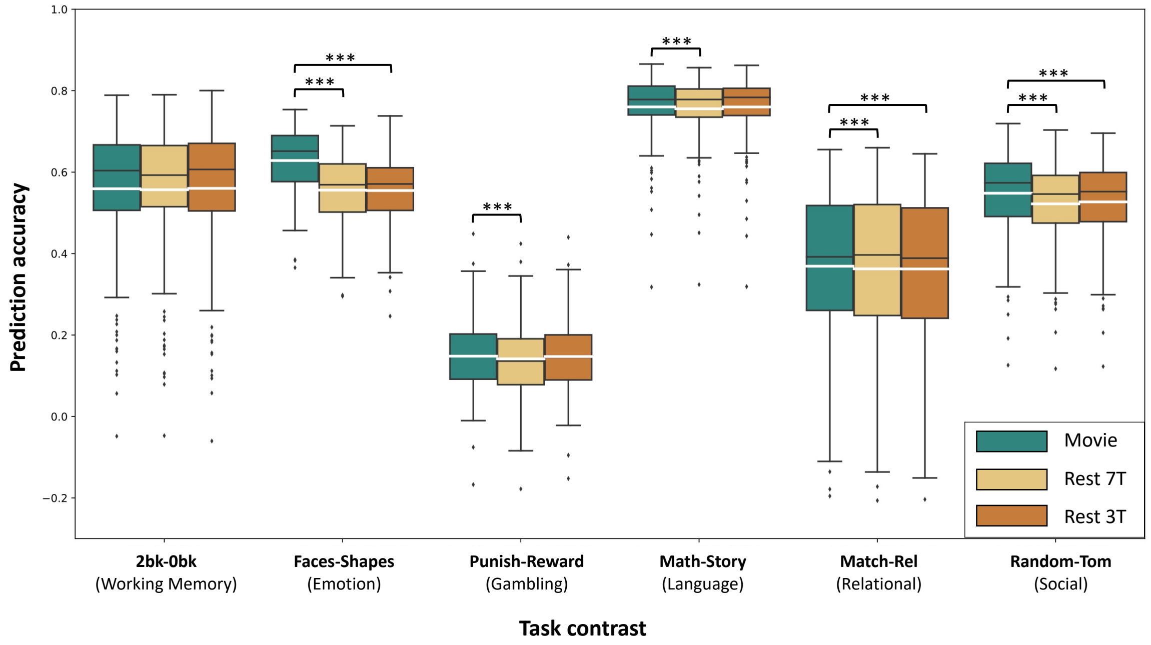


**Supplementary Figure S2. Movie-watching-derived *connTask* maps are more accurate than resting-state-derived maps.** This figure is similar to Figure 2 in the main test but shows task-contrasts between two conditions. For each task-contrast and each input type (movie-watching, 3T resting-state and 7T resting-state), boxplots portray the distribution of prediction accuracies. Prediction accuracy for each participant was calculated as the Pearson correlation between *connTask* and actual task-induced activation maps. Black lines represent the median, white lines represent the mean. Differences in *connTask* accuracy between input types and within each task-contrast were tested using a permutation test and corrected using Bonferroni correction for multiple comparisons. ***$p < 0.001$

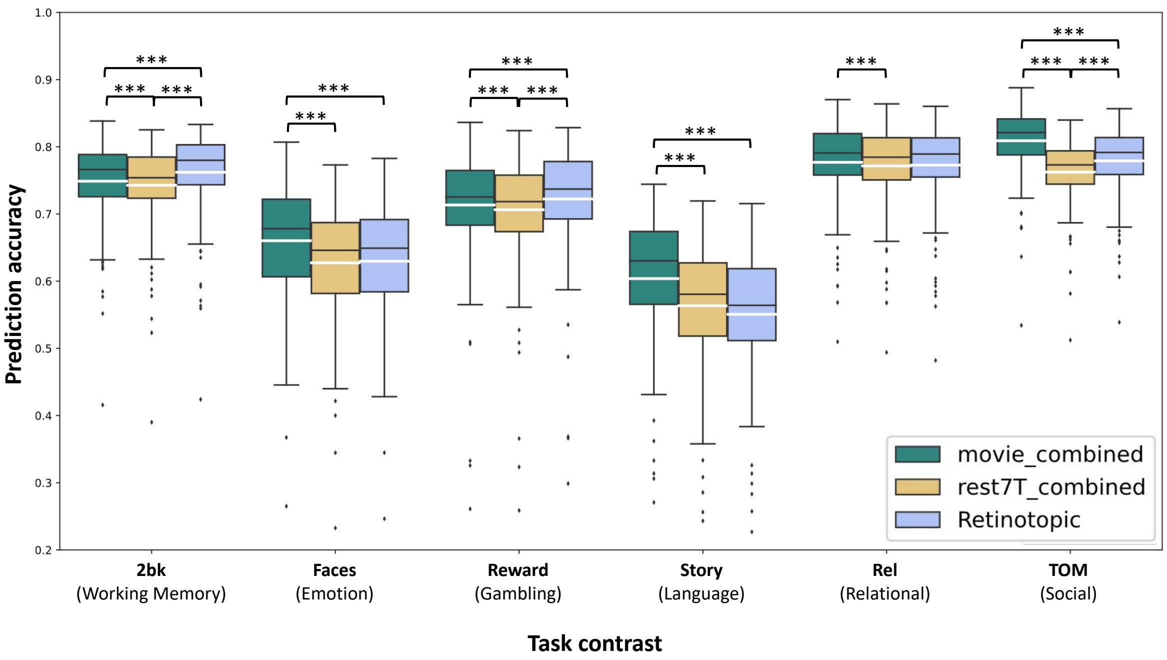


**Supplementary Figure S3. Movie-watching-derived *connTask* maps are more accurate than retinotopy-derived maps.** For each task-contrast and each input type (movie-watching, 7T resting-state and retinotopy), boxplots portray the distribution of prediction accuracies. Prediction accuracy for each participant was calculated as the Pearson correlation between *connTask* and actual task-induced activation maps. Black lines represent the median, white lines represent the mean. Differences in *connTask* accuracy between input types and within each task-contrast were tested using a permutation test and corrected using Bonferroni correction for multiple comparisons. ***$p < 0.001$

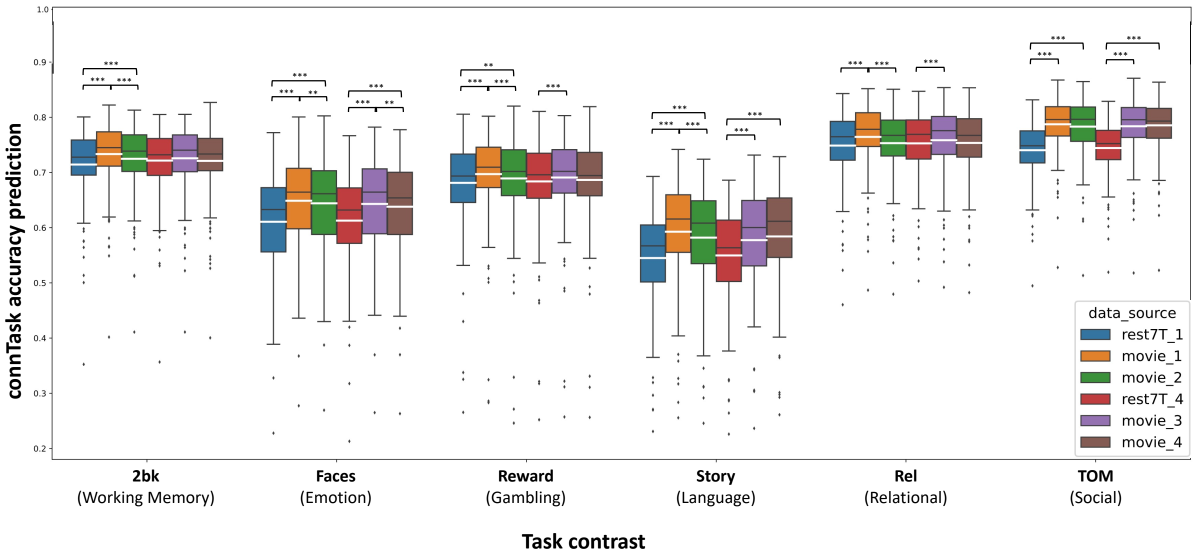


**Supplementary Figure S4. Accuracy of task-activity predictions derived from each movie-watching and resting-state run separately.** For each task-contrast and each input type, boxplots portray the distribution of prediction accuracies. Prediction accuracy for each participant was calculated as the Pearson correlation between *connTask* and actual task-induced activation maps. Black lines represent the median, white lines represent the mean. Differences in *connTask* accuracy between input types and within each task-contrast were tested using a permutation test and corrected using Bonferroni correction for multiple comparisons. ** $p<0.001$, *** $p<0.0001$

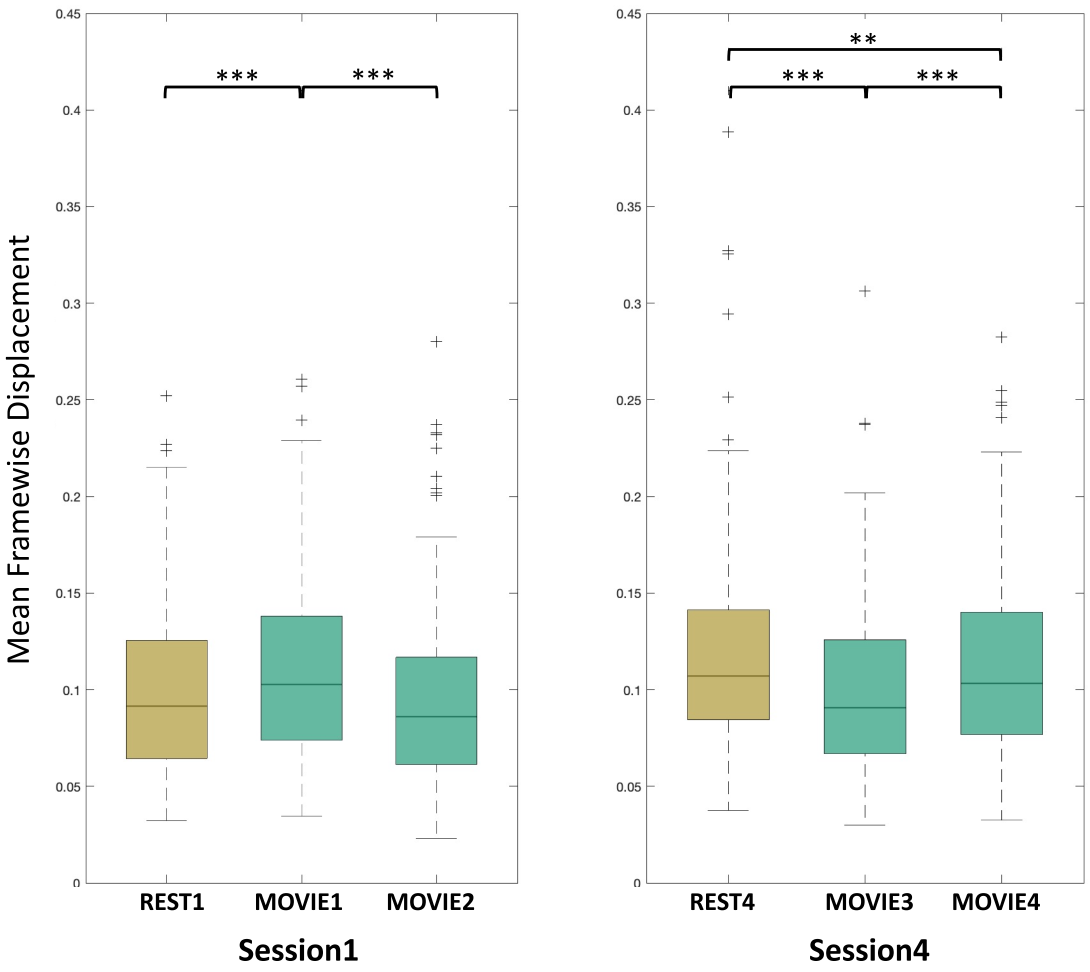


**Supplementary Figure S5. Differences in head motion in resting-state vs. movie-watching.** Mean framewise displacement was compared between resting-state and movie-watching runs within each session. In Session 1, larger head motion was observed while watching MOVIE1 compared to resting-state. In Session 2, larger head motion was observed during resting-state than while watching MOVIE3 and MOVIE4. ** $p<0.001$, *** $p<0.0001$

**Supplementary Table S1.** P-values and effect sizes of the difference in task-activity prediction accuracy between CC and HO movie-watching scans. The highlighted contrasts are the ones presented in Figure 4 in the main text and in Supplementary Figure S4.

| Task contrast | p-value | Cohen's D |
|---|---|---|
| Language_MATH | 0.006 | 0.19448 |
| Language_STORY | 0.066 | 0.12089 |
| Language_MATH_STORY | 0.0002 | 0.26428 |
| WM_2BK | 0.0002 | 0.33358 |
| WM_0BK | 0.0002 | 0.40779 |
| WM_2BK_0BK | 0.0002 | 0.2928 |
| Relational_MATCH | 0.0002 | 0.52446 |
| Relational_REL | 0.0002 | 0.48962 |
| Relational_MATCH_REL | 0.192 | 0.067835 |
| Gambling_PUNISH | 0.0002 | 0.29859 |
| Gambling_REWARD | 0.0002 | 0.3202 |
| Gambling_PUNISH_REWARD | 0.0486 | 0.13704 |
| Emotion_FACES | 0.0002 | 0.33505 |
| Emotion_SHAPES | 0.0002 | 0.58701 |
| Emotion_FACES_SHAPES | 0.9914 | 0.19223 |
| Social_RANDOM | 0.3296 | 0.039231 |
| Social_TOM | 0.1696 | 0.080448 |
| Social_RANDOM_TOM | 0.1206 | 0.094496 |
